# A deep learning-based drug repurposing screening and validation for anti-SARS-CoV-2 compounds by targeting the cell entry mechanism

**DOI:** 10.1101/2023.06.03.543589

**Authors:** Yingjia Yao, Yunhan Zhang, Zexu Li, Zhisong Chen, Xiaofeng Wang, Zihan Li, Li Yu, Xiaolong Cheng, Wei Li, Wen-Jie Jiang, Hua-Jun Wu, Zezhong Feng, Jinfu Sun, Teng Fei

## Abstract

The recent outbreak of Corona Virus Disease 2019 (COVID-19) caused by severe acute respiratory syndrome coronavirus 2 (SARS-CoV-2) has been a severe threat to the global public health and economy, however, effective drugs to treat COVID-19 are still lacking. Here, we employ a deep learning-based drug repositioning strategy to systematically screen potential anti-SARS-CoV-2 drug candidates that target the cell entry mechanism of SARS-CoV-2 virus from 2,635 FDA-approved drugs and 1,062 active ingredients from Traditional Chinese Medicine herbs. *In silico* molecular docking analysis validates the interactions between the top compounds and host receptors or viral spike proteins. Using a SARS-CoV-2 pseudovirus system, we further identify several drug candidates including Fostamatinib, Linagliptin, Lysergol and Sophoridine that can effectively block the cell entry of SARS-CoV-2 variants into human lung cells even at a nanomolar scale. These efforts not only illuminate the feasibility of applying deep learning-based drug repositioning for antiviral agents by targeting a specified mechanism, but also provide a valuable resource of promising drug candidates or lead compounds to treat COVID-19.

## 1. Introduction

Since the end of 2019, the severe respiratory coronavirus disease 2019 (COVID-19) caused by the infection of severe acute respiratory syndrome coronavirus 2 (SARS-CoV-2) has become a worldwide pandemic, leading to severely detrimental consequences to human health and society [1]. SARS-CoV-2 is an enveloped single stranded RNA virus with around 30 kb genome encoding four major structural proteins, including spike (S), nucleocapsid (N), membrane (M) and envelope proteins (E) [2]. The viral spike glycoprotein is the key molecule that mediates membrane fusion and entry into host cells. Human angiotensin-converting enzyme 2 (ACE2) is regarded as one of the master receptors for S protein to initiate cell entry process of SARS-CoV-2. Other membrane associated proteins such as furin-like proteases, transmembrane protease, serine 2 (TMPRSS2) and cathepsin L also play important roles as auxiliary receptors to facilitate viral entry into target cells [3]. Recent studies using genome-wide CRISPR screen or detailed mechanistic characterization further expand the collection of host factors that are required for SARS-CoV-2 cell entry or infection [4-7].

In addition to vaccination, developing effective drugs are necessary solutions to control COVID-19 [8]. Compared to *de novo* drug development, drug repositioning that repurposes “old drugs” to treat “new diseases” represents a more practical approach to identify safe and effective therapeutic agents against COVID-19 in a timely and cost-efficient manner. Actually, except for neutralizing antibodies, most of the current drugs approved or under clinical trials for the treatment of COVID-19 are derived from different drug repositioning strategies [8,9]. For instance, Remdesivir was originally synthesized as a nucleoside inhibitor of the respiratory syncytial virus. Ritonavir-boosted Nirmatrelvir (also known as Paxlovid) was initially developed as a protease inhibitor for SARS-CoV virus [8]. Thus, it is worthwhile to explore additional or complementary drug repurposing strategies for anti-SARS-CoV-2 agents. We previously developed a computational framework that integrates independent machine learning algorithms for predicting drug-target interaction (DTI) to efficiently repurpose existing marketed drugs and natural compounds in a high throughput manner against a given disease or mechanism by appropriately defining the target gene set [10,11]. Given the urgent and long-lasting detrimental effects of COVID-19, it is tentative to apply these approaches to continually explore repurposed drug candidates for anti-SARS-CoV-2 purpose.

Here we aim to identify potential antiviral drugs against SARS-CoV-2 by targeting the cell entry mechanism which is critical for the viral infection. By *in silico* analysis and experimental exploration, we firstly determined the expression pattern of a cohort of important membrane associated receptors or cofactors for SARS-CoV-2 cell entry in different human tissues and cell lines. Using key cell entry factors as the target gene set, we performed drug repositioning from thousands of FDA-approved drugs and natural compounds embedded in Traditional Chinese Medicine herbs with three independent deep learning algorithms. The top drug candidates exhibited significant interactions with the cell entry machinery as revealed by molecular docking analysis. Aided by a SARS-CoV-2 pesudovirus system, we successfully validated several compounds, such as FDA-approved drugs Fostomaticlib and Linagliptin as well as natural compounds Lysergol and Sophoridine, that can effectively inhibit SARS-CoV-2 variant entry into human lung cells. Collectively, these results provide valuable resources and lead compounds to develop anti-SARS-CoV-2 drugs targeting a defined cell entry mechanism.

## 2. Materials and Methods

### 2.1. Cell culture

Human A375, A549, HCT116, HEK293FT and Huh7 cells were cultured in DMEM medium (BIOTECH, REF: SC102-01), and SH-SY5Y cells were cultured in DMEM/F12 medium (BIOTECH, REF: SC103-01) supplemented with 10% fetal bovine serum (ExCell Bio, REF: FCS500) and 1% penicillin/streptomycin (Solarbio, Cat. No.: P1400) in a humid incubator at 37°C with 5% CO_2_.

### 2.2. Tested drug information

Chloroquine (CAS: 54-05-7), Fostamatinib (CAS: 901119-35-5), Linagliptin (CAS: 668270-12-0), Venetoclax (CAS: 1257044-40-8), Crizotinib (CAS: 877399-52-5), Lysergol (CAS: 602-85-7), and Sophoridine (CAS: 6882-68-4) were purchased from Shanghai Yuanye Bio-Technology company.

### 2.3. Quantitative reverse-transcription PCR (RT-qPCR)

RNA from cell samples were extracted with UNIQ-10 Column Trizol Total RNA Isolation Kit (Sangon Biotech, REF: B511321-0100), and 1 μg RNA per sample was reverse transcribed with RevertAid Reverse Transcription kit according to the manufacturer’s instructions (Thermo Fisher Scientific, REF: 4368814). Target transcript quantification was performed by real-time quantitative PCR with a reaction system comprising 2x UltraSYBR Mixture (CWBIO, REF: CW2601M), cDNA fraction and 10 μM forward / reverse primers. Fold-change was analyzed relative to human *RPS28* level using the ΔΔCt method. The primer sequences are listed in Supplementary Table 1.

**Table 1.** A list of top 5 repurposed FDA-approved drugs from each deep learning method.

| **Drug name** | **CAS no.** | **Typical indication** |
| --- | --- | --- |
| Alogliptin<sup>[c]</sup> | 850649-61-5 | Hyperglycemia in patients with type 2 diabetes mellitus |
| Bosutinib<sup>[a]</sup> | 380843-75-4 | Chronic, accelerated, or blast phase Philadelphia chromosome-positive chronic myelogenous leukemia |
| Crizotinib<sup>[a]</sup> | 877399-52-5 | Metastatic non-small cell lung cancer |
| Fostamatinib<sup>[c]</sup> | 901119-35-5 | Chronic immune thrombocytopenia |
| Linagliptin<sup>[c]</sup> | 668270-12-0 | Hyperglycemia in patients with type 2 diabetes mellitus |
| Neratinib<sup>[a]</sup> | 698387-09-6 | Breast cancer |
| Nintedanib<sup>[b]</sup> | 656247-17-5 | Idiopathic pulmonary fibrosis, systemic sclerosis-associated interstitial lung disease |
| Saxagliptin<sup>[c]</sup> | 361442-04-8 | Type 2 diabetes mellitus |
| Sitagliptin<sup>[c]</sup> | 486460-32-6 | Type 2 diabetes mellitus |
| Sunitinib<sup>[b]</sup> | 557795-19-4 | Renal cell carcinoma |
| Temsirolimus<sup>[a]</sup> | 162635-04-3 | Breast cancer |
| Venetoclax<sup>[a]</sup> | 1257044-40-8 | Chronic lymphocytic leukemia |
| Vinflunine<sup>[b]</sup> | 162652-95-1 | Advanced or metastatic transitional cell carcinoma of the urothelial tract |
| Vinorelbine<sup>[b]</sup> | 71486-22-1 | Metastatic non-small cell lung carcinoma |
| Voxilaprevir<sup>[b]</sup> | 1535212-07-7 | Hepatitis C infection |

[a] Drugs from DeepCPI method.
[b] Drugs from DeepPurpose method.
[c] Drugs from DTINet method.

### 2.4. Protein-protein interaction network analysis

The membrane associated cell entry-related proteins were imported into STRING database (Version 11.5, https://cn.string-db.org) to construct the protein-protein interaction network (PPI) with 0.15 as the minimally required interaction score, and the PPI network was visualized and analyzed using Cytoscape software (Version 3.9.1). Each diamond represents one protein, and each line of connections between the proteins represents the interaction relationship.

### 2.5. Drug repositioning for cell entry inhibitors

We described the detailed drug repositioning protocol in our previously studies[10,11]. To specifically target cell entry mechanism of SARS-CoV-2, here we used a cohort of reported cell entry factors and viral S proteins (Supplementary Table 2) as the target gene set. The drug library for repositioning contains 2635 FDA-approved drugs (DrugBank database, Version 5.1.7, released 2020-07-02; https://www.drugbank.ca) and 1062 active ingredients or natural compounds (Traditional Chinese Medicine Systems Pharmacology (TCMSP) online database, Version 2.3, released 2014-05-31; https://tcmspw.com/tcmsp.php). Three independent deep learning algorithms (DeepCPI[12], DTINet[13] and DeepPurpose[14]) were employed to calculate each drug-target interaction score and the drug candidates were finally ranked by the sum of drug-target interaction (DTI) score of all the target genes for each algorithm. The complete results of drug repurposing are deposited in Supplementary Dataset 1 and Supplementary Dataset 2.

**Table 2.** A list of top 5 repurposed natural compounds from each deep learning method.

| Compound Name | TCMSP ID<sup>[c]</sup> | PubChem Compound CID<sup>[d]</sup> | TCM |
| --- | --- | --- | --- |
| Atropine<sup>[a]</sup> | MOL002219 | 174174 | *Lycii Cortex* |
| Celabenzine<sup>[b]</sup> | MOL005314 | 163053085 | *Panax Ginseng C. A. Mey.* |
| Celafurine<sup>[b]</sup> | MOL003208 | 15346742 | *Tripterygii Radix* |
| Costaclavine<sup>[a]</sup> | MOL008145 | 160462 | *Ricini Semen* |
| Dauricumine<sup>[b]</sup> | MOL012904 | 11741544 | *Menispermii Rhizoma* |
| Lysergol<sup>[a]</sup> | MOL005261 | 14987 | *Semen Pharbitidis* |
| Solanocapsine<sup>[a]</sup> | MOL007356 | 73419 | *Solanum Nigrum Linn.* |
| Sophocarpine<sup>[a]</sup> | MOL003627 | 115269 | *Fructussophorae,*<br>*Sophorae Flavescentis Radix,*<br>*Sophorae Tonkinensis Radix Et*<br>*Rhizome* |
| Triptolide<sup>[b]</sup> | MOL003187 | 107985 | *Tripterygii Radix* |
| Triptonide<sup>[b]</sup> | MOL003244 | 65411 | *Tripterygii Radix* |

[a] Drugs from DeepCPI method.
[b] Drugs from DeepPurpose method.
[c] The data from TCMSP database (https://tcmsp-e.com/tcmsp.php).
[d] The data from PubChem database (https://pubchem.ncbi.nlm.nih.gov).

### 2.6. Molecular docking analysis

The 3D chemical structure of drugs were downloaded as SDF files from PubChem database (https://pubchem.ncbi.nlm.nih.gov), and the crystal structure of proteins were downloaded from RCSB PDB database (https://www.rcsb.org). The small molecules and proteins were then processed and optimized to generate PDBQT format files. The grid box was set to contain the entire protein region. The docking of drugs and proteins was performed with the genetic algorithm mode using AutoDock (Version 1.5.7). The docking results of the best possible conformations with the lowest energy were displayed by PyMOL software (Version 2.1) and the docking scores are listed in Supplementary Dataset 3.

### 2.7. Generation of SARS-CoV-2 pseudoviruses

We optimized the codon of SARS-CoV-2 spike (S) proteins for improved expression in human cells using GenSmart™ Codon Optimization Tool (GenScript, https://www.genscript.com), and obtained the S gene fragment of Wuhan-Hu-1 strain (GenBank: MN908947) and Omicron strain (GenBank: OW996240; B.1.1.529) by gene synthesis (GENEWIZ). To establish a lentivirus-based SARS-CoV-2 pseudovirus system, the vesicular stomatitis virus G envelope protein (VSV-G) element in pMD2.G plasmid was replaced by the corresponding spike of SARS-CoV-2 variants via molecular cloning between SapI and PmlI restriction enzyme sites. A lentivector pHAGE-EF1α-luciferase-BSD was used as the cargo transfer to deliver luciferase-expressing cassette in SARS-CoV-2 spike-pseudotyped viruses. We co-transfected pMD2.G-Wuhan-Hu-1 spike or pMD2.G-Omicron spike together with pCMVR8.74 and pHAGE-EF1α-luciferase-BSD plasmids into HEK293FT cells using Neofect™ DNA transfection reagent to produce the pseudoviruses. The supernatant was harvested after 48 h post transfection, centrifuged at 3000 rpm for 10 min to remove cellular debris and stored at - 80°C before use.

### 2.8. Luciferase assay

To determine the cell entry capability of SARS-CoV-2 pseudoviruses, the expression of pseudovirus-delivered luciferase in target cells were quantified. The human lung cancer A549 cells with stably ectopic expression of ACE2 were employed as target cells and seeded in 96-well plates at a density of 3 x 10^3^ cells/well. The cells were pre-treated with corresponding tested drugs for 12 h before pseudovirus infection and continually cultured for 48 h in the presence of tested compound and pseudoviruses. Then, 100 μL lysis buffer was added to each well of cells, and the luciferase activity was measured at λ = 560 nm in sextuplicate using a microplate reader (BioTek Synergy HTX).

### 2.9. Statistical analysis

All the experiments were independently performed for at least three times. GraphPad Prism 9.0 was used for statistical analysis. All data are expressed as means ± SD. Statistical significance of differences between two groups was analyzed with unpaired two-tailed t-test, and ^*^*p* < 0.05, ^**^*p* < 0.01 and ^***^*p* < 0.001 were considered statistically significant.

## 3. Results

### 3.1. The expression profiles of SARS-CoV-2 cell entry factors

The cell entry of SARS-CoV-2 relies on the intricate interplay between the viral spike glycoprotein and membrane-associated receptors as well as other auxiliary factors in host cells. In addition to well-characterized ACE2, furin, TMPRSS2 and cathepsin L, recent studies have identified more factors for SARS-CoV-2 cell entry in multiple cell lineages and models. We catalogued those important membrane associated cell entry factors by literature mining (Supplementary Table 2). By *in silico* analysis with RNA-seq data retrieved from NCBI Gene database and Cancer Cell Line Encyclopedia (CCLE) database, we firstly examined the RNA expression profiles of these genes across multiple human tissues and corresponding tissue-derived cell lines. As shown in Fig. 1A and 1B, these host cell entry factors displayed varied expression as quantified by RNA-seq read count in different human organ tissues including brain, colon, kidney, liver, lung, and skin, suggesting a tissue-specific preference for host factors engaged during SARS-CoV-2 cell entry. Consistent with previous reports [3], ACE2 showed higher expression in colon and kidney tissues (Fig. 1A). Although lung is a major target organ, the expression of ACE2 is primarily restricted in type II alveolar cells and upper bronchial epithelial cells, but not the whole lung [3]. In contrast, *CD147* and *GPR78* exhibited uniformly strong expression in many tissues, indicating their general and prevalent roles for SARS-CoV-2 infection. Notably, the expression pattern in tissues was generally consistent with that in corresponding tissue-derived cell lines (Fig. 1A and 1B), supporting a high quality of these expression data.

**Fig. 1.**
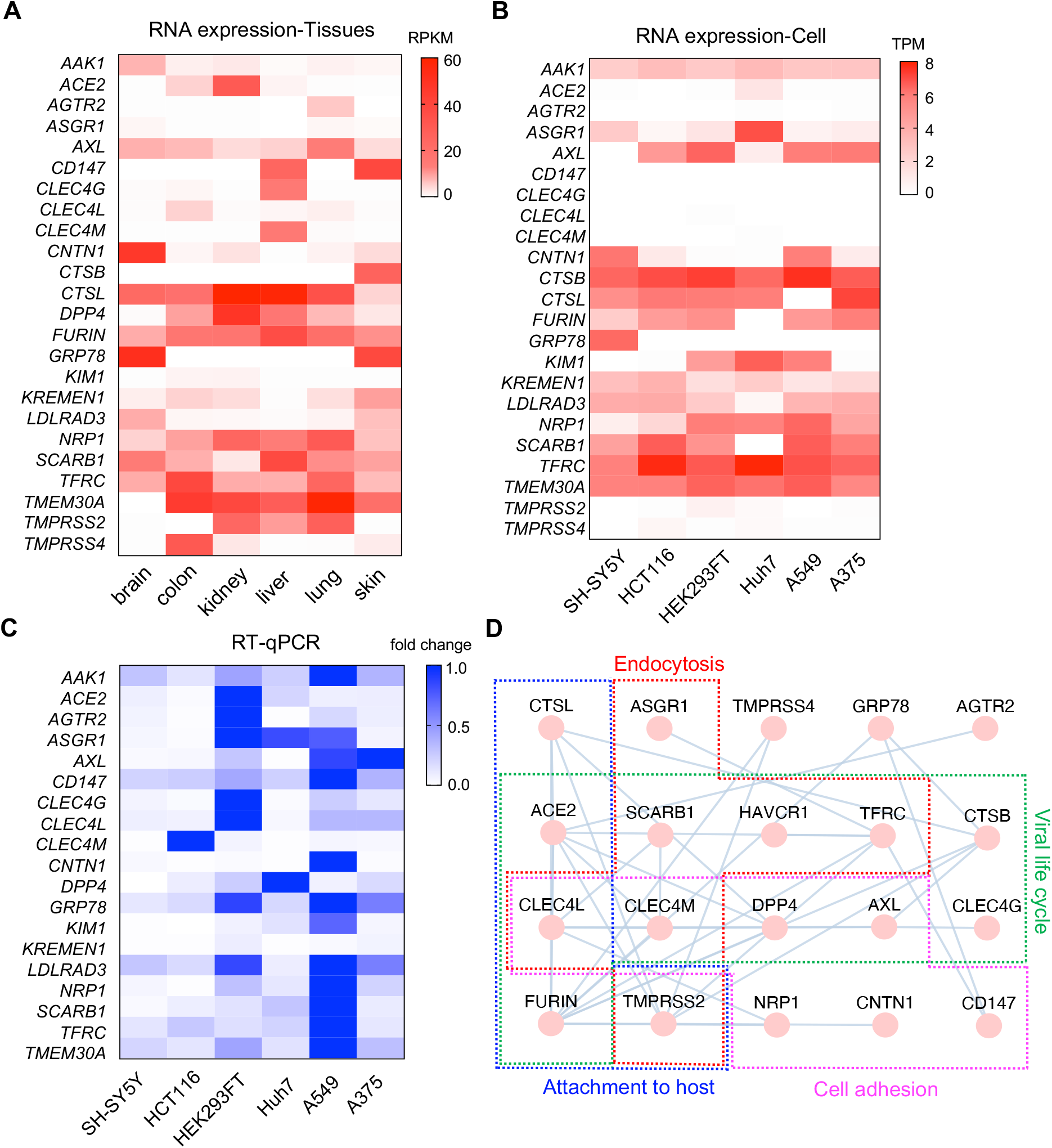
The expression profiles of SARS-CoV-2 cell entry factors. **A**. Heatmap showing the expressions of indicated SARS-CoV-2 cell entry genes across 6 different tissues in human with RPKM value from NCBI RNA-seq data. **B**. Heatmap showing the expressions of indicated genes in corresponding cell lines with TPM value from CCLE RNA-seq data. **C**. RT-qPCR validation of relative expression of indicated genes across different human cell lines. **D**. Protein-protein interaction network diagram showing the interactions and functional processes of indicated cell entry factors.

To further evaluate the expression levels of these cell entry genes and better select appropriate cell models for viral entry test, we extracted RNA from these cell lines and detected the relative RNA expression of indicated cell entry genes by quantitative reverse-transcription PCR (RT-qPCR). Unlike RPKM or TPM value in RNA-seq which enables a relative comparison of expression between different genes, the Ct value in qPCR may vary when using different primer pair even for the same gene, thus only allowing relative comparison between cell lines rather than across genes. For each gene, we set the relatively highest expression with a fold change value of 1.0 and other samples were standardized accordingly. As shown in Fig. 1C, we found that lung A549 cells expressed the highest levels of most cell entry factors, followed by HEK293FT cells. This pattern was generally consistent with the results obtained from *in silico* analysis (Fig. 1B). These data not only help to explain the differential tissue tropism of SARS-CoV-2 infection, but also suggest A549 cells among others as one of the most suitable cell models to explore cell entry of SARS-CoV-2. Moreover, we found that these cell entry factors have extensive interactions between each other and are functionally involved in critical steps of virus entry into target cells (Fig. 1D). These results prompt us to explore potential drugs that target these cell entry factors to block SARS-CoV-2 infection and alleviate COVID-19 symptoms.

### 3.2. High throughput drug repositioning to screen SARS-CoV-2 cell entry inhibitors

To efficiently identify potential anti-SARS-CoV-2 drugs specifically against the cell entry mechanism, we employed a deep learning-based drug repositioning strategy (Fig. 2A). In addition to host cell entry factors, we also included several major SARS-CoV-2 spike protein variants (Wuhan-Hu-1, Delta and Omicron) into the target gene set for drug repositioning. The drug library for repurposing includes 2,635 FDA-approved drugs and 1,062 active ingredients from TCM herbs. Three independent and computationally efficient algorithms DeepCPI, DTINet and DeepPurpose with great performance among others to date were chosen to calculate DTI score for each drug-target pair [12-14] (Fig. 2A). We finally ranked those repurposed drug hits for anti-SARS-CoV-2 cell entry according to their sum scores of potential interactions with each gene among the interrogated target gene set (Supplementary Dataset 1 and Supplementary Dataset 2). Notably, only two methods DeepCPI and DeepPurpose were applied for repositioning natural compounds since DTINet requires special drug information that is missing for natural compounds. Although the three algorithms employed various principles to calculate DTI score, we still observed some overlap of repurposed hits between different deep learning methods among the top 100 approved drugs or natural compounds (Fig. 2B). We reasoned that the results from different repositioning methods are complementary to each other and thus we highlighted the top 5 hits from each algorithm for approved drugs (Table 1) or natural compounds (Table 2), respectively.

**Fig. 2.**
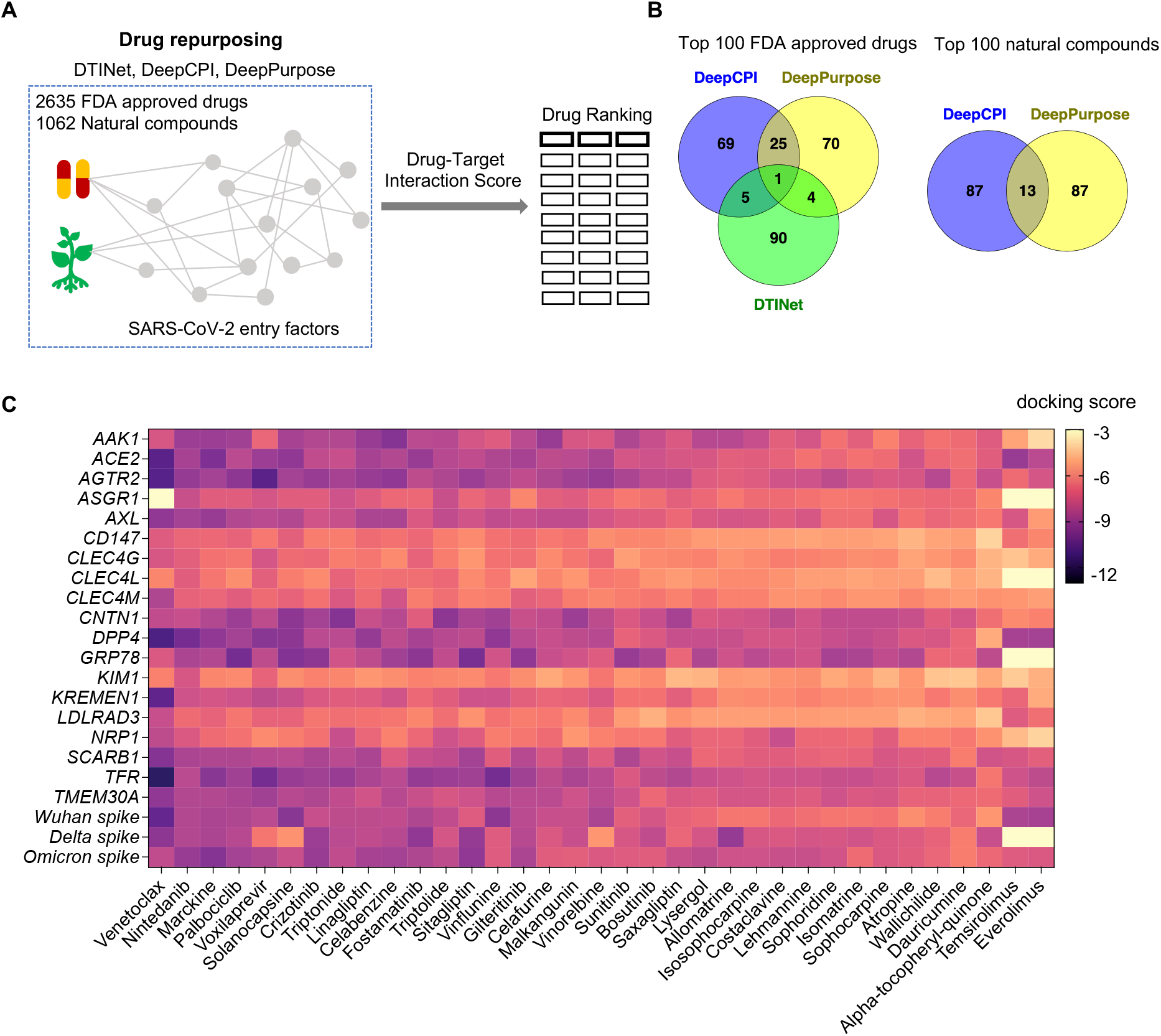
High throughput drug repositioning for anti-SARS-CoV-2 drugs by targeting cell entry factors. **A**. Strategic diagram of drug repositioning in this study. **B**. Venn diagram of the top repurposed anti-SARS-CoV-2 drug hits by different deep learning algorithms. **C**. The heatmap showing the score of molecular docking analysis for each indicated drug-target pair.

To systemically determine the potential interactions between each drug candidate and its target proteins, we performed a series of *in silico* structure-based molecular docking analysis for the top drug candidates and targeted cell entry factors. The matrix of docking score or predicted binding free energy was presented as a heatmap with a lower score indicating a stronger interacting potential and vice versa (Fig. 2C; Supplementary Dataset 3). We found that these top drug candidates exhibited a broad but differential capability to interact with multiple cell entry proteins. Certain host proteins (for example, ACE2, AGTR2, DPP4, GRP78 and TFR etc.) seemed to be preferentially targeted by those top drug hits with generally lower docking scores, suggesting that they may serve as primary targets for those drug candidates. On the other hand, three viral spike variants including Wuhan-Hu-1, Delta and Omicron strains also showed strong interaction with a broad range of top drug hits, supporting the key functions of S protein in determining viral entry into target cells. Interestingly, despite a very high similarity of protein sequences between the three S variants, some drugs (for instance, Temsirolimus and Everolimus) have significantly varied interaction potential for these S variants, further indicating pivotal roles of key amino acids of viral variants in defining the cell entry mechanisms and drug responses.

### 3.3. Experimental validation of drug candidates by SARS-CoV-2 cell entry assay

We next sought to experimentally validate the effects of the top repurposed drug candidates on blocking SARS-CoV-2 cell entry. Due to the restricted biosafety requirement to operate authentic viruses, we instead adopted a well-established SARS-CoV-2 spike-pseudotyped lentivirus system which delivers a luciferase-expressing cassette as a relevant and convenient surrogate approach to study SARS-CoV-2 cell entry [15] (Fig. 3A). We packed pseudoviruses for both Wuhan-Hu-1 and Omicron SARS-CoV-2 variants and chose ACE2-expressing A549 cells as suitable target cells based on our above analysis. The intensity of luciferase activity was used as the readout for SARS-CoV-2 cell entry and infectivity (Fig. 3A). From the top drug candidates, we managed to obtain six compounds from commercially available sources (Materials and Methods) and proceeded to test their effects on the cell entry of SARS-CoV-2 pseudoviruses. Using chloroquine as a positive control for SARS-CoV-2 cell entry [16], we were able to observe four tested drugs (Fostamatinib, Linagliptin, Lysergol and Sophoridine) that exhibited a dose-dependent effect to block SARS-CoV-2 cell entry while other two drugs Venetoclax and Crizotinib did not have such effects (Fig. 3B and 3C; Supplementary Fig. 1). The inhibitory effect existed for both Wuhan-Hu-1 and Omicron variants with only a few differences of inhibition magnitude (Fig. 3B and 3C). Notably, for Linagliptin, Lysergol and Sophoridine, either compound can inhibit the infection of one or two SARS-CoV-2 variants up to 40% at a low 1 μM concentration, and the blocking effect manifested even at a nanomolar scale. Compared to these three compounds, the effect of Fostamatinib was relatively weaker in this model system (Fig. 3B and 3C). Furthermore, these effective compounds did not show obvious cell toxicity across tested concentration range (Fig. 3D). Molecular docking analysis showed strong interaction potential between these compounds and representative cell entry targets (Fig. 3E). These data suggest that Fostamatinib, Linagliptin, Lysergol and Sophoridine hold potential to serve as safe and potent cell entry inhibitors of SARS-CoV-2 to treat COVID-19.

**Fig. 3.**
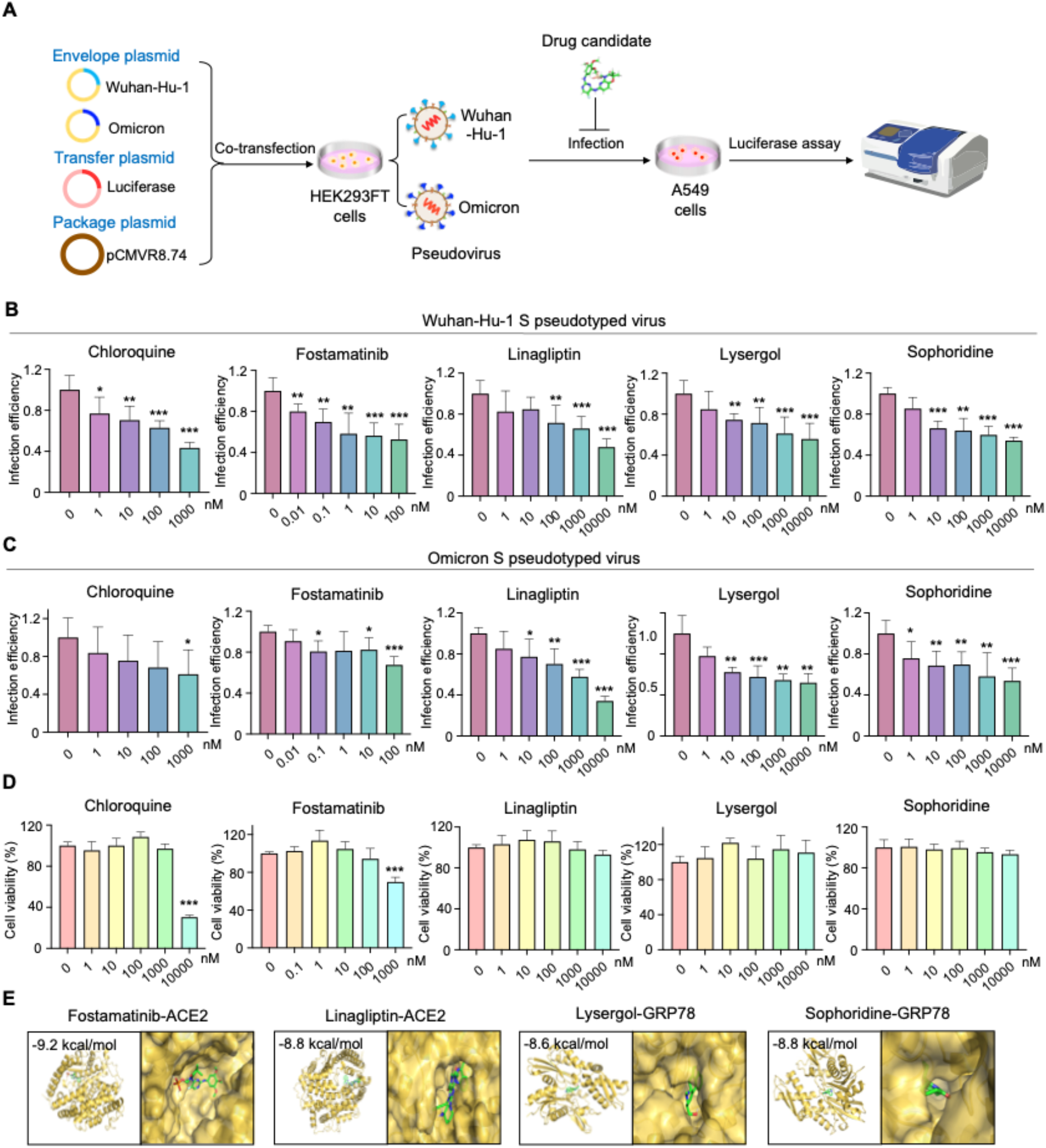
Experimental validation of selective repurposed drugs to inhibit SARS-CoV-2 pseudovirus cell entry. **A**. Schematic workflow of repurposed drug validation using SARS-CoV-2 pseudoviruses. **B-C**. Infection efficiency of Wuhan-Hu-1 S pseudotyped virus (**B**) and Omicron S pseudotyped virus (**C**) for ACE2-expressing A549 cells in the presence of indicated doses of tested compounds measured by luciferase assay. **D**. Cell toxicity of indicated doses of tested compounds for ACE2-expressing A549 cells measured by MTT-based cell viability assay. **E**. The representative docking results for indicated drugs and targeted proteins.

## 4. Discussion

Developing effective anti-SARS-CoV-2 therapeutics is still an urgently unmet medical need in the context of devastating COVID-19 threat. In this study, we performed a deep learning-based drug repositioning screening to identify potential SARS-CoV-2 inhibitors that target cell entry mechanism during viral infection. Among the top ranked hits, we were able to validate several approved drugs and natural compounds that can effectively block the entry of SARS-CoV-2 pseudoviruses into human lung cells even a low nanomolar concentration.

Compared to previous drug repositioning studies for anti-SARS-CoV-2 agents, the novelty and significance of our work here mainly lie in the following aspects: (1) We primarily target the cell entry factors in the host as host-directed therapeutics may better deal with the rapidly mutated viral variants to avoid frequent target alteration or immune escape for virus-acting agents; (2) Current host-acting drugs for COVID-19 are mainly immunomodulators to control infection-associated immune or inflammatory responses, but very few host-directed drug candidates were proposed to target the appealing cell entry process of SARS-CoV-2 [8]; (3) Unlike previous studies that usually target a single (e.g., ACE2 or spike) or unknown (direct compound screening) gene during drug repositioning [17-20], we take multiple host cell entry factors as well as viral S proteins as a whole target gene set into consideration which may provide more accurate and balanced information to better select drug candidates; (4) Three independent deep learning algorithms are applied for improved prediction power during drug repurposing; (5) We employ more relevant human lung A549 cells rather than the widely used monkey kidney Vero E6 cells for SARS-CoV-2 cell entry assay which may help to better translate the results into human clinics; (6) In addition to FDA-approved drugs, we also include thousands of natural compounds in TCM herbs for drug repurposing, which provides another useful pool of lead compounds or active ingredients to develop anti-SARS-CoV-2 drugs.

Among the validated drug candidates, Fostamatinib is a spleen tyrosine kinase (Syk) inhibitor approved for the treatment of chronic immune thrombocytopenia. It also has inhibitory activity for Janus kinase (JAK) and therefore is suggested as an immunomodulator to improve COVID-19-associated immune dysregulation [8]. Our data here suggest that Fostamatinib might have another therapeutic action by directly inhibiting SARS-CoV-2 cell entry and thus could be applied in early stages of COVID-19 disease rather than just immunomodulator administrated in late stages for hospitalized patients. Linagliptin is an inhibitor of Dipeptidyl Peptidase-4 (DPP4) and used to control blood sugar in adults with type 2 diabetes [21]. Another drug repurposing study predicted Linagliptin as a potential inhibitor of viral cysteine protease by molecular docking [22], while our work proposed its action in blocking viral cell entry with experimental evidence. Although Linagliptin did not show clear effects in patients hospitalized with both COVID-19 and diabetes [23,24], our results indicate that general populations with early infection of SARS-CoV-2 may benefit from this oral drug. Sophoridine is a bioactive alkaloid found in many TCM herbs. Given that other similar alkaloids such as Matrine and Oxysophoridine have been implicated in the treatment of COVID-19 [25,26], it is reasonable for Sophoridine to exert an anti-SARS-CoV-2 function. Lysergol is also an alkaloid found in some fungi and TCM herbs with certain anti-bacterial activity [27]. It is interesting to observe its anti-SARS-CoV-2 roles, and further work is needed to fully characterize this potential lead compound.

In summary, this study demonstrates the power of applying multiple deep learning methods and specifically targeting the cell entry mechanism to screen antiviral agents by drug repositioning. And more importantly, we identified several approved drugs and natural compounds as promising therapeutic agents for the prevention and treatment of COVID-19.

## Acknowledgements

This project was funded by the Science and Technology Plan Project of Liaoning Province (2022JH2/20100001), the 111 Project (B16009), and Key Laboratory of Bioresource Research and Development of Liaoning Province (2022JH13/10200026) to T.F.; and the Fundamental Research Funds for the Central Universities (N2120006; N2224002-13) to Y.Y.

## Appendix A. Supplementary data

Supplementary data to this article can be found online.

## References

[1] J. Li, S. Lai, G.F. Gao, W. Shi, The emergence, genomic diversity and global spread of SARS-CoV 2, Nature 600 (2021) 408–418. 10.1038/s41586-021-04188-6.

[2] M.M. Lamers, B.L. Haagmans, SARS-CoV-2 pathogenesis, Nat Rev Microbiol 20 (2022) 270–284. 10.1038/s41579-022-00713-0.

[3] C.B. Jackson, M. Farzan, B. Chen, H. Choe, Mechanisms of SARS-CoV-2 entry into cells, Nat Rev Mol Cell Biol 23 (2022) 3–20. 10.1038/s41580-021-00418-x.

[4] J. Wei, M.M. Alfajaro, P.C. DeWeirdt, R.E. Hanna, W.J. Lu-Culligan, W.L. Cai, M.S. Strine, S.M. Zhang, V.R. Graziano, C.O. Schmitz, J.S. Chen, M.C. Mankowski, R.B. Filler, N.G. Ravindra, V. Gasque, F.J. de Miguel, A. Patil, H. Chen, K.Y. Oguntuyo, L. Abriola, Y.V. Surovtseva, R.C. Orchard, B. Lee, B.D. Lindenbach, K. Politi, D. van Dijk, C. Kadoch, M.D. Simon, Q. Yan, J.G. Doench, C.B. Wilen, Genome-wide CRISPR Screens Reveal Host Factors Critical for SARS-CoV-2 Infection, Cell 184 (2021) 76–91 e13. 10.1016/j.cell.2020.10.028.

[5] Y. Zhu, F. Feng, G. Hu, Y. Wang, Y. Yu, Y. Zhu, W. Xu, X. Cai, Z. Sun, W. Han, R. Ye, D. Qu, Q. Ding, X. Huang, H. Chen, W. Xu, Y. Xie, Q. Cai, Z. Yuan, R. Zhang, A genome-wide CRISPR screen identifies host factors that regulate SARS-CoV-2 entry, Nat Commun 12 (2021) 961. 10.1038/s41467021-21213-4.

[6] A. Rebendenne, P. Roy, B. Bonaventure, A.L. Chaves Valadao, L. Desmarets, M. Arnaud-Arnould, Y. Rouille, M. Tauziet, D. Giovannini, J. Touhami, Y. Lee, P. DeWeirdt, M. Hegde, S. Urbach, K.E. Koulali, F.G. de Gracia, J. McKellar, J. Dubuisson, M. Wencker, S. Belouzard, O. Moncorge, J.G. Doench, C. Goujon, Bidirectional genome-wide CRISPR screens reveal host factors regulating SARS CoV-2, MERS-CoV and seasonal HCoVs, Nat Genet 54 (2022) 1090–1102. 10.1038/s41588-02201110-2.

[7] S.B. Biering, S.A. Sarnik, E. Wang, J.R. Zengel, S.R. Leist, A. Schafer, V. Sathyan, P. Hawkins, K. Okuda, C. Tau, A.R. Jangid, C.V. Duffy, J. Wei, R.C. Gilmore, M.M. Alfajaro, M.S. Strine, X. Nguyenla, E. Van Dis, C. Catamura, L.H. Yamashiro, J.A. Belk, A. Begeman, J.C. Stark, D.J. Shon, D.M. Fox, S. Ezzatpour, E. Huang, N. Olegario, A. Rustagi, A.S. Volmer, A. Livraghi-Butrico, E. Wehri, R.R. Behringer, D.J. Cheon, J. Schaletzky, H.C. Aguilar, A.S. Puschnik, B. Button, B.A. Pinsky, C.A. Blish, R.S. Baric, W.K. O’Neal, C.R. Bertozzi, C.B. Wilen, R.C. Boucher, J.E. Carette, S.A. Stanley, E. Harris, S. Konermann, P.D. Hsu, Genome-wide bidirectional CRISPR screens identify mucins as host factors modulating SARS-CoV-2 infection, Nat Genet 54 (2022) 1078–1089. 10.1038/s41588-022-01131-x.

[8] G. Li, R. Hilgenfeld, R. Whitley, E. De Clercq, Therapeutic strategies for COVID-19: progress and lessons learned, Nat Rev Drug Discov (2023) 1–27. 10.1038/s41573-023-00672-y.

[9] J. Yin, C. Li, C. Ye, Z. Ruan, Y. Liang, Y. Li, J. Wu, Z. Luo, Advances in the development of therapeutic strategies against COVID-19 and perspectives in the drug design for emerging SARS-CoV 2 variants, Comput Struct Biotechnol J 20 (2022) 824–837. 10.1016/j.csbj.2022.01.026.

[10] Z. Li, Y. Yao, X. Cheng, Q. Chen, W. Zhao, S. Ma, Z. Li, H. Zhou, W. Li, T. Fei, A computational framework of host-based drug repositioning for broad-spectrum antivirals against RNA viruses, iScience 24 (2021) 102148. 10.1016/j.isci.2021.102148.

[11] Z. Li, Y. Yao, X. Cheng, W. Li, T. Fei, An in silico drug repositioning workflow for host-based antivirals, STAR Protoc 2 (2021) 100653. 10.1016/j.xpro.2021.100653.

[12] F. Wan, Y. Zhu, H. Hu, A. Dai, X. Cai, L. Chen, H. Gong, T. Xia, D. Yang, M.W. Wang, J. Zeng, DeepCPI: A Deep Learning-based Framework for Large-scale in silico Drug Screening, Genomics Proteomics Bioinformatics 17 (2019) 478–495. 10.1016/j.gpb.2019.04.003.

[13] Y. Luo, X. Zhao, J. Zhou, J. Yang, Y. Zhang, W. Kuang, J. Peng, L. Chen, J. Zeng, A network integration approach for drug-target interaction prediction and computational drug repositioning from heterogeneous information, Nat Commun 8 (2017) 573. 10.1038/s41467-017-00680-8.

[14] K. Huang, T. Fu, L.M. Glass, M. Zitnik, C. Xiao, J. Sun, DeepPurpose: a deep learning library for drug-target interaction prediction, Bioinformatics 36 (2021) 5545–5547. 10.1093/bioinformatics/btaa1005.

[15] M. Xu, M. Pradhan, K. Gorshkov, J.D. Petersen, M. Shen, H. Guo, W. Zhu, C. Klumpp-Thomas, S. Michael, M. Itkin, Z. Itkin, M.R. Straus, J. Zimmerberg, W. Zheng, G.R. Whittaker, C.Z. Chen, A high throughput screening assay for inhibitors of SARS-CoV-2 pseudotyped particle entry, SLAS Discov 27 (2022) 86–94. 10.1016/j.slasd.2021.12.005.

[16] M. Wang, R. Cao, L. Zhang, X. Yang, J. Liu, M. Xu, Z. Shi, Z. Hu, W. Zhong, G. Xiao, Remdesivir and chloroquine effectively inhibit the recently emerged novel coronavirus (2019-nCoV) in vitro, Cell Res 30 (2020) 269–271. 10.1038/s41422-020-0282-0.

[17] Q. Tong, G. Liu, X. Sang, X. Zhu, X. Fu, C. Dou, Y. Jian, J. Zhang, S. Zou, G. Zhang, X. Du, D. Liu, S. Qi, W. Cheng, Y. Tian, X. Fu, Targeting RNA G-quadruplex with repurposed drugs blocks SARS-CoV-2 entry, PLoS Pathog 19 (2023) e1011131. 10.1371/journal.ppat.1011131.

[18] Y. Chen, Y. Wu, S. Chen, Q. Zhan, D. Wu, C. Yang, X. He, M. Qiu, N. Zhang, Z. Li, Y. Guo, M. Wen, L. Lu, C. Ma, J. Guo, W. Xu, X. Li, L. Li, S. Jiang, X. Pan, S. Liu, S. Tan, Sertraline Is an Effective SARS-CoV-2 Entry Inhibitor Targeting the Spike Protein, J Virol 96 (2022) e0124522. 10.1128/jvi.01245-22.

[19] R.R.R. Duarte, D.C. Copertino, Jr., L.P. Iniguez, J.L. Marston, Y. Bram, Y. Han, R.E. Schwartz, S. Chen, D.F. Nixon, T.R. Powell, Identifying FDA-approved drugs with multimodal properties against COVID-19 using a data-driven approach and a lung organoid model of SARS-CoV-2 entry, Mol Med 27 (2021) 105. 10.1186/s10020-021-00356-6.

[20] L. Yang, R.J. Pei, H. Li, X.N. Ma, Y. Zhou, F.H. Zhu, P.L. He, W. Tang, Y.C. Zhang, J. Xiong, S.Q. Xiao, X.K. Tong, B. Zhang, J.P. Zuo, Identification of SARS-CoV-2 entry inhibitors among already approved drugs, Acta Pharmacol Sin 42 (2021) 1347–1353. 10.1038/s41401-020-00556-6.

[21] K. McKeage, Linagliptin: an update of its use in patients with type 2 diabetes mellitus, Drugs 74 (2014) 1927–1946. 10.1007/s40265-014-0308-3.

[22] P.P.N. Rao, A.T. Pham, A. Shakeri, A. El Shatshat, Y. Zhao, R.C. Karuturi, A.A. Hefny, Drug Repurposing: Dipeptidyl Peptidase IV (DPP4) Inhibitors as Potential Agents to Treat SARS-CoV-2 (2019-nCoV) Infection, Pharmaceuticals (Basel) 14 (2021) 10.3390/ph14010044.

[23] R. Guardado-Mendoza, M.A. Garcia-Magana, L.J. Martinez-Navarro, H.E. Macias-Cervantes, R. Aguilar-Guerrero, E.L. Suarez-Perez, A. Aguilar-Garcia, Effect of linagliptin plus insulin in comparison to insulin alone on metabolic control and prognosis in hospitalized patients with SARS-CoV-2 infection, Sci Rep 12 (2022) 536. 10.1038/s41598-021-04511-1.

[24] R. Abuhasira, I. Ayalon-Dangur, N. Zaslavsky, R. Koren, M. Keller, D. Dicker, A. Grossman, A Randomized Clinical Trial of Linagliptin vs. Standard of Care in Patients Hospitalized With Diabetes and COVID-19, Front Endocrinol (Lausanne) 12 (2021) 794382. 10.3389/fendo.2021.794382.

[25] Y.N. Zhang, Q.Y. Zhang, X.D. Li, J. Xiong, S.Q. Xiao, Z. Wang, Z.R. Zhang, C.L. Deng, X.L. Yang, H.P. Wei, Z.M. Yuan, H.Q. Ye, B. Zhang, Gemcitabine, lycorine and oxysophoridine inhibit novel coronavirus (SARS-CoV-2) in cell culture, Emerg Microbes Infect 9 (2020) 1170–1173. 10.1080/22221751.2020.1772676.

[26] M.W. Yang, F. Chen, D.J. Zhu, J.Z. Li, J.L. Zhu, W. Zeng, S.L. Qu, Y. Zhang, [Clinical efficacy of Matrine and Sodium Chloride Injection in treatment of 40 cases of COVID-19], Zhongguo Zhong Yao Za Zhi 45 (2020) 2221–2231. 10.19540/j.cnki.cjcmm.20200323.501.

[27] A. Maurya, G.R. Dwivedi, M.P. Darokar, S.K. Srivastava, Antibacterial and synergy of clavine alkaloid lysergol and its derivatives against nalidixic acid-resistant Escherichia coli, Chem Biol Drug Des 81 (2013) 484–490. 10.1111/cbdd.12103.

