## Supplementary Information for "A deep learning-based drug repurposing screening and validation for anti-SARS-CoV-2 compounds by targeting the cell entry mechanism"

**This file includes:**

**Supplementary Figure 1.** Experimental validation of repurposed compounds by SARS-CoV-2 pseudovirus cell entry assay.
**Supplementary Table 1.** Primer sequences for RT-qPCR.
**Supplementary Table 2.** A list of important membrane associated cell entry factors for SARS-CoV-2.
**Supplementary References**

**Other Supplementary Materials for this manuscript include the following (separate file):**

**Supplementary Dataset 1.** Drug repurposing results for FDA-approved drugs. (attached dataset)
**Supplementary Dataset 2.** Drug repurposing results for natural compounds. (attached dataset)
**Supplementary Dataset 3.** Molecular docking results for top drug candidates and key cell entry factors. (attached dataset)

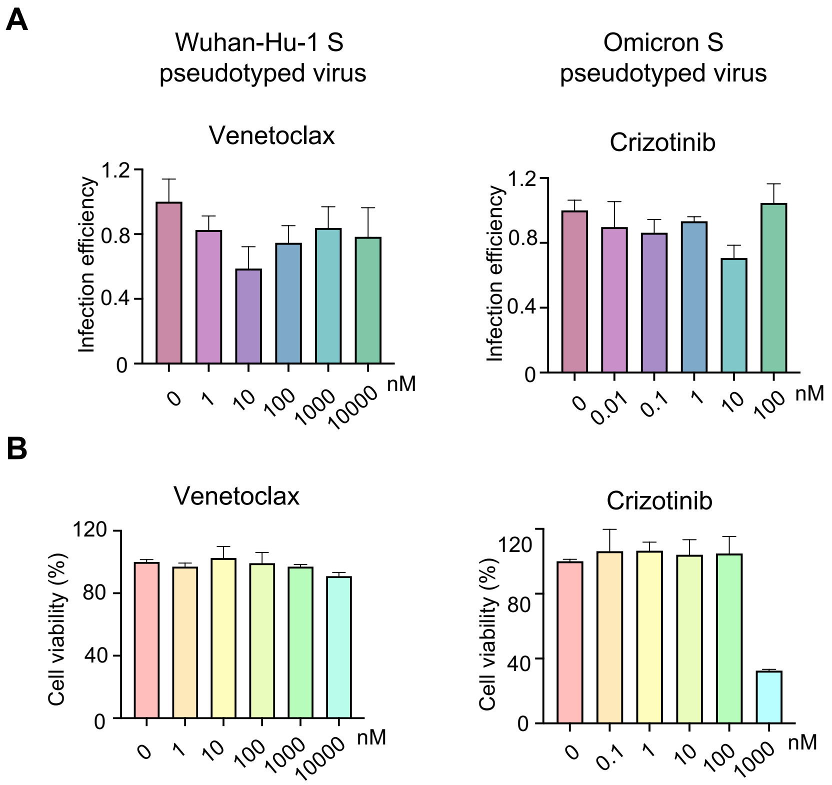


**Supplementary Fig. 1.** Experimental validation of repurposed compounds by SARS-CoV-2 pseudovirus cell entry assay. **A.** Infection efficiency of Wuhan-Hu-1 S pseudotyped virus and Omicron S pseudotyped virus for ACE2-expressing A549 cells in the presence of indicated doses of tested compounds measured by luciferase assay. **B.** Cell toxicity of indicated doses of tested compounds for ACE2-expressing A549 cells measured by MTT-based cell viability assay.

**Supplementary Table 1.** Primer sequences for RT-qPCR.

| Gene | Forward-sequence | Reverse-sequence |
|---|---|---|
| AAK1 | AGTGGCTACATCGGAAGAGTC | AGGCACATTTCATCCCATTGC |
| ACE2 | ACAGTCCACACTTGCCCAAAT | TGAGAGCACTGAAGACCCATT |
| AGTR2 | CTGGCAACAATGAGTCTACCTT | GCCACAGCGAGGTTGAAGAT |
| ASGR1 (CLEC4H1) | ATGAAGTCGCTAGAGTCCCAG | CAGGTCAGACACGAACTGCTT |
| AXL (UFO) | GTGGGCAACCCAGGGAATATC | GTACTGTCCCGTGTCGGAAAG |
| CD147 (BSG) | GACGACCAGTGGGGAGAGTA | GGCCTTGTCCTCAGAGTCAG |
| CLEC4G | AGTCCTTTGGGCTGTGATTCT | AGGCGTTTGTCCTCAGCAG |
| CLEC4L (CD209) | AATGGCTGGAACGACGACAAA | CAGGAGGCTGCGGACTTTTT |
| CLEC4M (CD209L) | ACTTCATGTCTAACTCCCAGCG | ATTCCGCACAGTCTTCATTCC |
| CNTN1 | CAGCCCTTTCCCGGTTTACAA | TGCTTCTGACCATCCCGTAGT |
| DPP4 (CD26) | GGGTCACATGGTCACCAGTG | TCTGTGTCGTTAAATTGGGCATA |
| GRP78 (HSPA5) | GAAAGAAGGTTACCCATGCAGT | CAGGCCATAAGCAATAGCAGC |
| KIM1 (HAVCR1) | TGGCAGATTCTGTAGCTGGTT | AGAGAACATGAGCCTCTATTCCA |
| KREMEN1 | CCCGAGTGTTTCACAGCCAAT | GGATGCTGGAAAGTCTCGTTC |
| LDLRAD3 | CAATGAGTGCAACATACCAGGC | ACTCTTGTCGAAGCAGTCAGG |
| NRP1 | ACCCAAGTGAAAAATGCGAATG | CCTCCAAATCGAAGTGAGGGTT |
| SCARB1 (SR-B1) | CCTATCCCCTTCTATCTCTCCG | GGATGTTGGGCATGACGATGT |
| TFR (TFRC) | ACCATTGTCATATACCCGGTTCA | CAATAGCCCAAGTAGCCAATCAT |
| TMEM30A | CAAAACCATCGTCGTTACGTGA | GTTGGCAATAGCTCCACAAGG |
| RPS28 | CGATCCATCATCCGCAATG | AGCCAAGCTCAGCGCAAC |

**Supplementary Table 2.** A list of important membrane associated cell entry factors for SARS-CoV-2.

| Protein | Description | Tissue specificity[a] | Biological process or molecular function[b] | Disorder[c] | Reference |
|---|---|---|---|---|---|
| AAK1 | AP2 Associated Kinase 1 | Parathyroid, gland | Endocytosis, Serine/threonine-protein kinase | Rabies, Hepatitis C Virus, Ebola Hemorrhagic Fever | [1] |
| ACE2 | Angiotensin I converting enzyme 2 | Gallbladder, Intestine, Kidney | Host-virus interaction, Carboxypeptidase | Cardiovascular System Disease, Heart Disease, Covid-19, Severe Acute Respiratory Syndrome, etc. | [2] |
| AGTR2 | Angiotensin II Receptor Type 2 | Endometrium, Lung, Smooth Muscle | G-protein coupled receptor | Obstructive Nephropathy, Covid-19, Vesicoureteral Reflux, Ureteral Obstruction, etc. | [3] |
| ASGR1 | Asialoglycoprotein receptor 1 | Liver | Endocytosis, Host-virus interaction | Hepatitis, Acute Gonococcal Cervicitis, Autoimmune Hepatitis, etc. | [4] |
| AXL | AXL Receptor Tyrosine Kinase | Low tissue specificity | Differentiation, Host-virus interaction, Host cell receptor for virus entry | Hypogonadotropic Hypogonadism With or Without Anosmia, NK-Cell Enteropathy | [5] |
| CD147 | (Basigin) Ok Blood Group | Low tissue specificity | Host-virus interaction, Angiogenesis | Severe Acute Respiratory Syndrome, Ameloblastoma, Melanoma, cancer, etc. | [6] |

| | | | | | |
|---|---|---|---|---|---|
| CLEC4G | C-Type Lectin Domain Family 4 Member G | Adipose tissue, Liver, Lymphoid tissue | Host cell receptor for virus entry, Host-virus interaction | Lymphocytic Choriomeningitis, Japanese Encephalitis, Encephalitis, Argentine Hemorrhagic Fever, etc. | [7] |
| CLEC4L | C-type lectin domain family 4 member L | Adipose tissue, Placenta | Host cell receptor for virus entry, Adaptive immunity, Cell adhesion | Dengue Virus, Human Immunodeficiency Virus Type 1, Rift Valley Fever, Japanese Encephalitis, etc. | [8] |
| CLEC4M | C-type lectin domain family 4 member M | Liver, Lymphoid tissue | Adaptive immunity, Endocytosis, Host-virus interaction, Innate immunity | West Nile Virus, Japanese Encephalitis, Hepatitis C, Encephalitis, etc. | [9] |
| CNTN1 | Contactin-1 | Brain | Cell adhesion | Polyradiculopathy, Autoimmune Neuropathy, Myopathy, Miller Fisher Syndrome, etc. | [10] |
| CTSB | Cathepsin B | General granular cytoplasmic expression | Hydrolase, Protease, Thiol protease | Squalene Synthase Deficiency, Tropical Calcific Pancreatitis, Occlusion of Gallbladder, Bone Giant Cell Tumor. | [11] |
| CTSL | Cathepsin L | Lysosomal expression in several tissues | Hydrolase, Protease, Thiol protease, Host-virus interaction | Covid-19, Middle East Respiratory Syndrome, Benign Meningioma, Severe Acute Respiratory Syndrome, etc. | [12] |
| DPP4 | Dipeptidyl Peptidase 4 | Intestine, Parathyroid gland, Placenta, Prostate | Cell adhesion, Aminopeptidase | Middle East Respiratory Syndrome, Hyperglycemia, Nasopharyngitis, Covid-19, etc. | [9] |

| | | | | | |
|---|---|---|---|---|---|
| FURIN | FURIN | Liver, Salivary gland | Heparin-binding, Hydrolase, Protease, Serine protease, Host-virus interaction | Diphtheria, Anthrax Disease, Cerebral Amyloid Angiopathy, Mumps, etc. | [2] |
| GRP78 | Heat Shock Protein Family A (Hsp70) Member 5 | Low tissue specificity | Host-virus interaction, Hydrolase | Mucormycosis, Japanese Encephalitis, Borna Disease, Prion Disease, etc. | [13] |
| KIM1 | Hepatitis A virus cellular receptor 1 | Intestine, Kidney | Host-virus interaction | Hepatitis, Dengue Virus, Obstructive Nephropathy, Chikungunya, etc. | [14] |
| KREMEN1 | Kringle containing transmembrane protein 1 | Low tissue specificity | Wnt signaling pathway | Ectodermal Dysplasia, Hair/Tooth Type, Hyperostosis, Schopf-Schulz-Passarge Syndrome, etc. | [4] |
| LDLRAD3 | Low Density Lipoprotein Receptor Class A Domain Containing 3 | Low tissue specificity | Host-virus interaction, Receptor | Venezuelan Equine Encephalitis, Postinflammatory Pulmonary Fibrosis, Western Equine Encephalitis. | [15] |
| NRP1 | Neuropilin 1 | Low tissue specificity | Host-virus interaction, Angiogenesis, Developmental protein | Covid-19, Pancreatic Cancer, Cerebral Arteriopathy, Neuroma, etc. | [16] |
| SCARB1 | Scavenger Receptor Class B Member 1 | Adrenal gland | Host cell receptor for virus entry, Host-virus interaction | Hepatitis C, Hyperalphalipoproteinemia, Platelet Glycoprotein Iv Deficiency, Cholelithiasis, etc. | [17] |
| TFR | Transferrin Receptor | Bone marrow | Host cell receptor for virus entry, Endocytosis | Immunodeficiency, Combined Immunodeficiency. | [18] |

| | | | | | |
|---|---|---|---|---|---|
| TMEM30A | Transmembrane Protein 30A | Low tissue specificity | Lipid transport | Cholestasis, Benign Recurrent Intrahepatic, X-Linked Nephrolithiasis Type I. | [19] |
| TMPRSS2 | Transmembrane serine protease 2 | Prostate, Stomach | Host-virus interaction | Covid-19, Influenza, Middle East Respiratory Syndrome, Severe Acute Respiratory Syndrome, etc. | [2] |
| TMPRSS4 | Transmembrane serine protease 4 | Liver, Salivary gland | Hydrolase, Protease, Serine protease, Host-virus interaction | Deafness, Covid-19, Lynch Syndrome. | [20] |

[a] The data from Human Protein Atlas database (https://www.proteinatlas.org).

[b] The data from UniProt database (https://www.uniprot.org).

[c] The data from GeneCards database (https://www.genecards.org).

### Supplementary References


1. Ghamry, H.I., et al., *Evaluating the ability of some natural phenolic acids to target the main protease and AAK1 in SARS COV-2*. Sci Rep, 2023. **13**(1): p. 7357.
2. Jackson, C.B., et al., *Mechanisms of SARS-CoV-2 entry into cells*. Nat Rev Mol Cell Biol, 2022. **23**(1): p. 3-20.
3. Cui, C., et al., *AGTR2, One Possible Novel Key Gene for the Entry of SARS-CoV-2 Into Human Cells*. IEEE/ACM Trans Comput Biol Bioinform, 2021. **18**(4): p. 1230-1233.
4. Gu, Y., et al., *Receptome profiling identifies KREMEN1 and ASGR1 as alternative functional receptors of SARS-CoV-2*. Cell Res, 2022. **32**(1): p. 24-37.
5. Wang, S., et al., *AXL is a candidate receptor for SARS-CoV-2 that promotes infection of pulmonary and bronchial epithelial cells*. Cell Res, 2021. **31**(2): p. 126-140.
6. Wang, K., et al., *CD147-spike protein is a novel route for SARS-CoV-2 infection to host cells*. Signal Transduct Target Ther, 2020. **5**(1): p. 283.
7. Hoffmann, D., et al., *Identification of lectin receptors for conserved SARS-CoV-2 glycosylation sites*. EMBO J, 2021. **40**(19): p. e108375.
8. Amraei, R., et al., *CD209L/L-SIGN and CD209/DC-SIGN Act as Receptors for SARS-CoV-2*. ACS Cent Sci, 2021. **7**(7): p. 1156-1165.
9. Zhang, G.F., et al., *Infectivity of pseudotyped SARS-CoV-2 variants of concern in different human cell types and inhibitory effects of recombinant spike protein and entry-related cellular factors*. J Med Virol, 2023. **95**(1): p. e28437.

10. Husain, B., et al., *Cell-based receptor discovery identifies host factors specifically targeted by the SARS CoV-2 spike*. Commun Biol, 2022. **5**(1): p. 788.
11. Gao, X., et al., *Genome-wide screening of SARS-CoV-2 infection-related genes based on the blood leukocytes sequencing data set of patients with COVID-19*. J Med Virol, 2021. **93**(9): p. 5544-5554.
12. Zhao, M.M., et al., *Cathepsin L plays a key role in SARS-CoV-2 infection in humans and humanized mice and is a promising target for new drug development*. Signal Transduct Target Ther, 2021. **6**(1): p. 134.
13. Carlos, A.J., et al., *The chaperone GRP78 is a host auxiliary factor for SARS-CoV-2 and GRP78 depleting antibody blocks viral entry and infection*. J Biol Chem, 2021. **296**: p. 100759.
14. Zhang, F., et al., *SARS-CoV-2 pseudovirus infectivity and expression of viral entry-related factors ACE2, TMPRSS2, Kim-1, and NRP-1 in human cells from the respiratory, urinary, digestive, reproductive, and immune systems*. J Med Virol, 2021. **93**(12): p. 6671-6685.
15. Zhu, S., et al., *Genome-wide CRISPR activation screen identifies candidate receptors for SARS-CoV-2 entry*. Sci China Life Sci, 2022. **65**(4): p. 701-717.
16. Cantuti-Castelvetri., L., et al., *Neuropilin-1 facilitates SARS-CoV-2 cell entry and infectivity*. Science, 2020. **370**: p. 856-860.
17. Wei, C., et al., *HDL-scavenger receptor B type 1 facilitates SARS-CoV-2 entry*. Nat Metab, 2020. **2**(12): p. 1391-1400.
18. Wang, X., et al., *Transferrin Receptor Protein 1 Is an Entry Factor for Rabies Virus*. J Virol, 2023. **97**(2): p. e0161222.
19. Alipoor, S.D. and M. Mirsaeidi, *SARS-CoV-2 cell entry beyond the ACE2 receptor*. Mol Biol Rep, 2022. **49**(11): p. 10715-10727.
20. KATOPOdIS., P., et al., *COVID-19 and SARS-CoV-2 host cell entry mediators: Expression profiling of TMRSS4 in health and disease*. International Journal of Molecular Medcine, 2021. **47**(64).
